## Supplementary Information for "Bilateral Human Laryngeal Motor Cortex in Perceptual Decision of Lexical Tone and Voicing of Consonant"

### Continua generation

##### Experiment 1

Continua were synthesized by Matlab R2016a and legacy-STRAIGHT. Each continuum included multiple steps of syllables, which varied in only one acoustic property (VOT or F0 contour). Two types of continua were used: consonant continua ranging from [t] to [t^h^] (“d” to “t” in Pinyin; note that the “d” in Pinyin is the unaspirated voiceless dental plosive, instead of the voiced [d]^1^), as well as tone continua ranging from high-level tone [55] to mid-rising tone [35] (tone1 to tone2). To generate a VOT continuum, implemented by manual code in Matlab, short segments (1/54 the length of natural VOT) of aspiration noise were stepwise appended between release-burst and the phonation onset of the unaspirated [t] until reaching [t^h^]. Also, to keep the authenticity of syllables, phonation periods of a VOT continuum were also morphed from [ti] to [t^h^i] using STRAIGHT synthesis algorithms, but this could hardly affect categorical perception since VOT is the most significant feature to distinguish these two consonants. To make an F0 continuum, a pair of syllables differing only in tone (i.e., one with high-level tone and another one mid-rising tone, but sharing the same VOT) were entered into STRAIGHT synthesis algorithms. Further, a 54×54-step matrix of tone–consonant continua was generated by orthogonal morphing. Hence, each syllable in the matrix represented a step in both tone and consonant continua.

5×5-step individualized continuum matrices were generated to fit participants’ ranges of perceptual ambiguity (steps in a continuum where participants were unable to determine). To estimate the ranges for tone and consonant continua, in a preliminary experiment, participants performed tone or consonant identification tasks with a 9×9-step matrix selected from the 54×54-step matrix, using the method of constant stimuli. In each trial, participants heard a syllable, and needed to identify to which tone/consonant category the syllable belonged, ignoring the other dimension. Edges of ranges were manually detected. Based on the estimated edges, individualized continua were morphed as was described above (see Supplementary Table 1 for acoustic properties of the individualized continua, Supplementary Audio 2 for F0 continuum, Supplementary Audio 3 for VOT continuum).

Before the preliminary experiment started, participants were familiarized with the four natural syllables by listening to each of them. Then, they finished a 20-trial four-alternative forced-choice practice task of syllable identification. A correct rate of more than 90% (i.e., > 18 in 20 trials) was the requirement for entry. Afterward, to ensure that participants understood the tasks, they finished a discrimination task to determine if the heard pair of syllables shared an identical tone/consonant category. The discrimination task was continued until participants discriminated all pairs correctly.

##### Experiment 2

The same syllables [ti55], [ti35], and [t^h^i55], corresponding to Mandarin syllables “堤dī” (riverbank), “敌dí” (enemy), and “梯tī” (ladder) from Experiment 1 were used in Experiment 2. Different from Experiment 1, only one 54-step tone continuum ([ti55] to [ti35]) and one 54-step consonant continuum ([ti55] to [t^h^i55]) were synthesized. Procedures of stimuli preprocessing and continuum morphing were identical to Experiment 1.

In Experiment 2, both ranges of perceptual ambiguity and SNR levels for noise masking were individualized. Participants first entered familiarization sessions identical to Experiment 1. To estimate individual ranges of perceptual ambiguity, a preliminary experiment similar to that in Experiment 1 was conducted, but participants performed tone and consonant identification tasks using tone and consonant continuum, respectively. Individualized SNR levels were estimated by the method of constant stimuli where participants performed tone and consonant identification tasks using unambiguous syllables ([ti55] and [ti35] for tone tasks, [ti55] and [t^h^i55] for consonant tasks) masked by SSN. SSN (identical to Experiment 1 except 5 kHz low-pass filtered) began and ended simultaneously with the syllable, with additional 10 ms linear rise-decay envelopes. For each task, the SNR level where participants reached around 85% of decision accuracy was defined as the individualized SNR level: -16dB to 2dB in tone tasks; -12dB to 12dB in consonant tasks.

### Behavioral data analysis

##### Preprocessing of the slope

For sham condition, slopes with negative values (< 0) or with PSE out of the continuum interval (< 1 or > 5) were eliminated as these parameters were mathematically invalid in the current context. For rTMS conditions, the “mathematically invalid” slopes could indicate that rTMS completely disrupted categorical perception. Therefore, instead of being deleted, these invalid slopes were replaced by the floor values in the corresponding sham blocks. To get the floor values, outliers (3SD from mean) were removed from the preprocessed data for each condition. Then, the block-wise floor values were derived from the formula F = exp(mean(log(D)) - 2*SD(log(D))), where D was the data while F was the corresponding floor value. The preprocessed sham slopes and the floor value-adjusted rTMS slopes were logarithmically transformed to adjust for positive skewness (See Supplementary Fig. 1 for a graphical illustration of this procedure).

##### Analysis of TMS effects

TMS effects were defined as differences between slopes in TMS blocks and the corresponding sham blocks, and were statistically tested using single-tailed one-sample permutation test in Matlab 2016a (if TMS - Sham < 0 or not). Compared with t-test, permutation test does not require the normality assumption of sample distribution, as distributions of slopes in some conditions were skewed as estimated the by Kolmogorov-Smirnov test of normality. To perform a permutation test, for each condition in TMS blocks, we randomly selected participants and permuted their slopes in the TMS block and sham block, contrasted TMS to sham, and calculated the mean value. This permutation–contrast–average process was repeated 10^5^ times and a permutation distribution of random mean differences was generated. P-value was calculated by ranking the real TMS effect among values in the permutation distribution: p = (ranking + 1) / (10^5^ +1). In addition, for Experiment 1, the relative down-regulation of rTMS upon the dLMC on categorical perception was estimated by comparing dLMC stimulation with TMC stimulation effects on slopes ((dLMC - Sham) - (TMC - Sham)) using single-tailed Wilcoxon signed-rank test in Matlab 2016a. For Experiment 2, the relative down-regulation of cTBS upon the dLMC compared with iTBS was assessed in similar ways ((cTBS - Sham) - (iTBS - Sham)). Here, we used Wilcoxon signed-rank test instead of permutation test because the different TMS conditions (dLMC and TMS stimulation in Experiment 1, and iTBS and cTBS stimulation in Experiment 2) shared the same sham condition, and it was unable to simultaneously permute the sham condition with two TMS conditions. To correct for multiple comparisons, FDR correction was used with p threshold = 0.05. In Experiment 1, a group of comparisons refers to one group of trials (ambiguous, half-ambiguous, and unambiguous) in each group of participants (stimulating left or right hemisphere). In Experiment 2, a group of comparisons refers to all conditions the group of participants underwent.

##### Competitions between consonant and tone perception

To explore interactions between consonant and tone perception, we used data from sham conditions in Experiment 1 (Experiment 1 Sham) as interference from TMS was excluded. Participants in left and right hemisphere stimulation groups were pooled together. Here, to extract slopes, for each block of tasks, trials were categorized into ambiguous (step 3), half-ambiguous (steps 2 and 4), and unambiguous (steps 1 and 5) conditions according to the unattended dimension of a matrix (Supplementary Fig. 3a). This allows us to test the main effects of ambiguity of the unattended feature as well as to quantify the task difficulty as a function of such ambiguity.

To confirm the reliability of potential effects, we also analyzed data from two other unpublished tests (Supplementary Experiment 1 & 2) conducted previously. Both tests used paradigms similar to this study, but in groups of participants that did not enter the current study.

22 right-handed young adults (13 females, mean age 21.32 years, SD = 2.16) participated in Supplementary Experiment 1. The group size is sufficient to detect medium to large effects as estimated by G*Power3. They had normal hearing and gave written informed consent prior to the experiment. They were paid ￥50 per hour for participation. The experiment was approved by the Ethics Committee of the Institute of Psychology, Chinese Academy of Sciences. Participants performed speeded tone/consonant categorical identification tasks with/without noise similar to those in Experiment 1. Stimuli came from a 7×7-steps tone (high-level tone to mid-rising tone) – consonant ([t] to [t^h^]) matrix of continua. One token was presented in each trial and each block contained 200 trials (more trials around PSE to guarantee curve fitting), to which different tokens were randomly assigned. Inter-trial interval (ITI) was on average 0.5s (0.4–0.6s, 10 ms step, uniformly distributed). For each block, trials were categorized into ambiguous (step 4), half-ambiguous (steps 2, 3, 5, and 6), and unambiguous (steps 1 and 7) conditions according to the unattended dimension of a matrix.

58 right-handed young adults (29 females, mean age = 21.31 years, SD = 2.93) participated in Supplementary Experiment 2. The group size is sufficient to detect medium to large effects as estimated by G*Power3. They had normal hearing and gave written informed consent prior to the experiment. They were paid ￥50 per hour for participation. The experiment was approved by the Ethics Committee of the Institute of Psychology, Chinese Academy of Sciences. Participants performed speeded tone and consonant categorical identification tasks similar to those in Experiment 1, but only in quiet. Stimuli came from a 5×5-step matrix of continua (high-level tone to mid-rising tone)–([t] to [t^h^]). Each block contained 39 trials, to which different tokens were randomly assigned. ITI was identical to that in Supplementary Experiment 1. For each block, trials were categorized into ambiguous (step 3), half-ambiguous (steps 2 and 4), and unambiguous (steps 1 and 5) conditions according to the unattended dimension of a matrix.

The fitting and preprocessing procedures for Experiment 1 Sham, Supplementary Experiment 1, and Supplementary Experiment 2 were identical to those in TMS experiments (Supplementary Fig. 1). Notably, curve fitting was performed separately for each ambiguity condition of trials, and block-level upper and lower asymptotes were calculated by unconstrained fit on data.

Slopes of psychometric curves in each block were pooled by averaging them across blocks (tone or consonant blocks for all three tests, and with/without noise mask for Experiment 1 Sham and Supplementary Experiment 1) separately for ambiguous, half-ambiguous, and unambiguous conditions of trials. For each test, the main effects of ambiguity of the unattended feature on the slopes were tested. To do so, we applied one-way repeated measures analysis of variance (rANOVA) to Experiment 1 Sham and Supplementary Experiment 1. For Supplementary Experiment 2, however, since the Kolmogorov-Smirnov test of normality showed that slopes in the unambiguous condition were not normally distributed, the Friedman test, a non-parametric alternative of rANOVA was used instead. Results showed that, main effects of ambiguity of the unattended feature were significant in all three tests (Experiment 1 Sham: *F*_2,74_ = 4.266, p = 0.018, partial η^2^ = 0.103; Supplementary Experiment 1: *F*_2,30_ = 5.001, p = 0.013, partial η^2^ = 0.250; Supplementary Experiment 2: χ^2^(2) = 6.259, p = 0.044, Kendall’s W = 0.058).

Moreover, we applied post-hoc contrast analyses to investigate how the ambiguity of the unattended feature affects tone and consonant categorical perception. Results showed that, for all three tests, slopes were lowered when the unattended feature was half-ambiguous. In Experiment 1 Sham, slopes from the half-ambiguous condition were lower than those from the ambiguous condition (Mean difference = -0.331, p = 0.022, Bonferroni corrected). In Supplementary Experiment 1, slopes from the half-ambiguous condition were lower than those from the ambiguous (Mean difference = -0.425, p = 0.042, Bonferroni corrected) and unambiguous (Mean difference = -0.403, p = 0.021, Bonferroni corrected) conditions. In Supplementary Experiment 2, slopes from the half-ambiguous condition were lower than those from the unambiguous condition (*Z* = -2.820, p = 0.014, Bonferroni corrected, Wilcoxon signed-rank test). Presumably, processing of the unattended feature of a syllable may occupy neural resources for the parallel computations of the attended feature^2^, and such competition was amplified when the unattended feature was half-ambiguous as participants had biased but uncertain perception.

### Functional localization experiment

##### Procedure

48 right-handed participants (26 females, mean age = 21.44, SD = 2.79) took part in the functional localization experiment of speech motor areas (i.e., dLMC and TMC). The group size is sufficient to detect medium to large effects as estimated by G*Power3. They gave written informed consent prior to the experiment. They were paid ￥100 per hour for participation. The experiment was approved by the Ethics Committee of the Institute of Psychology, Chinese Academy of Sciences. 10 participants also took part in Experiment 1 (N = 9) and Experiment 2 (N = 1). Before the experiment, T1-weighted anatomical images were acquired on a 3-Tesla Siemens Magnetom Trio scanner using a 20-channel head coil, using a magnetization-prepared rapid acquisition gradient echo (MPRAGE) sequence (TR = 2200 ms, TE = 3.49 ms, slices per slab = 192, field of view FOV = 256 mm, flip angle = 8°, spatial resolution = 1×1×1 mm), identical to Experiment 1 and 2. During the experiment, participants needed to articulate the voiceless [t] (only tongue moved) or pronounce the voiced [a] (only laryngeal vocal fold vibrated), to localize the TMC and the dLMC, respectively. The experiment was in a block design and was divided into 8 blocks, including 4 TMC and 4 LMC localization blocks. The order of blocks was counterbalanced across participants. A resting period with the same duration followed each block. Each block contained 8 trials, each lasting for 2s. To avoid breathing artifacts, before the beginning of each block, participants took a deep breath and held their breath until the block ended. In each trial, participants articulated or pronounced sounds following the visual cue (“Say AH” or “Say D”) projected to the screen. Functional images were acquired by a continuous multiband-accelerated echo-planar imaging sequence (multiband factor = 4, 40 slices, TR = 640 ms, TE = 30 ms, flip angle = 25°, FOV = 192 mm, voxel size = 3 × 3 × 3 mm). Blood oxygen level-dependent (BOLD) signals were preprocessed and analyzed at individual and group levels.

##### fMRI data analysis

Preprocessing of anatomical and functional images were implemented by fMRIprep, an fMRI preprocessing pipeline that incorporates multiple state-of-art software tools^3^.

T1-weighted (T1w) anatomical images were corrected for intensity non-uniformity and skull-stripped with ANTs 2.3.3^4^ (RRID:SCR_004757), and used as T1w-reference. Spatial normalization to standard space was performed through nonlinear registration with ANTs 2.3.3. The template used for spatial normalization was FSL’s MNI ICBM 152 non-linear 6th Generation Asymmetric Average Brain Stereotaxic Registration Model^5^ (RRID:SCR_002823; TemplateFlow ID: MNI152NLin6Asym).

To process functional images, fMRIprep was used to correct for susceptibility distortions. Co-registrations of the BOLD to anatomical references were performed by ANTs 2.3.3. Head-motion parameters were estimated by FSL 5.0.9^6^. Slice-timing correction was performed using AFNI 20160207^7^. The BOLD time-series were resampled and realigned to the MNI152NLin6Asym standard space. AFNI 20220424 was applied for spatial smoothing (Gaussian filter FWHM = 6.0 mm) and scaling the BOLD time-series to a mean of 100 to derive percent signal change.

Generalized linear model (GLM) analyses were conducted at the individual level by AFNI 20220424. The predicted time course of BOLD activation was modeled as a “box-car” function convolved with the canonical hemodynamic response function. Individual contrast maps were defined by comparing “AH” blocks to “D” blocks (dLMC: “AH” – “D”, TMC: “D” – “AH”) using paired t-test. Contrast maps were then subjected to group analysis where one-sample t-test (alternative hypothesis μ = 0) was used to find activated brain areas, which were thresholded by multiple comparison corrections (3dttest++ ClustSim program in AFNI 20220424, uncorrected p < 0.001, 10000 Monte Carlo simulations, smallest cluster size for “AH” – “D” = 239 voxels, “D” – “AH” = 164 voxels), and were masked by ROIs of bilateral precentral gyrus in automated anatomical labeling (AAL) templates^8^.

MNI coordinates of bilateral homologous dLMC and TMC were determined by visual inspection on the group map of activation. Criteria of target selection included: (1) bilateral homologous targets should share a similar degree of activation; (2) geometric middle points, instead of peaks of activation areas, were selected to ensure that magnetic stimulation covers the maximal volume of activations; (3) the dLMC and TMC targets should be spatially dissociated that TMS on either site would not affect the other (i.e., out of the 1.5 cm effective radius of the 70 mm figure-of-eight coil we used^9^). As for results, the spatial patterns of activations were similar to previous studies^10,11^, except that we did not find group-level significant activations in the ventral LMC. Whether the phonation area is located in the dorsal or the ventral or both of the LMC is still debated, but studies and models have emphasized the role of the dorsal LMC in human voice and pitch regulations^12^ (See Discussion for details). Thus, the current study only focuses on the dorsal LMC. The chosen MNI coordinates were: [±40, -5, 50] for dLMC and [±59, -3, 36] for TMC (Supplementary Fig. 4). Note that, although the two selected targets were proximal to each other, their spatial distance still exceeds the effective range of a figure-eight TMS coil^9^. Thus, transmission effects of stimulation can be ignored.

### Drift-diffusion model and reaction time analysis

##### Interactions between cTBS effects and stimulus ambiguity

To study whether cTBS effects on DDM parameters were affected by stimulus ambiguity (i.e., the step in a continuum), we built 8 regression models with various interaction items based on the full model that assumes cTBS affected all three parameters (the winning model, see Methods, Drift-diffusion model and reaction time analysis). These models recruit none (baseline), one, two, or all (full) of the three parameters, respectively. Note that the “baseline” model here is equal to the full model without interaction items. We used dummy treatment coding with the intercept set on sham conditions. Each model drew 2000 posterior samples, and discarded the first 20 samples as burn-in. Models are shown below (AMB: stimulus ambiguity).

**Baseline model:**

*a* ~ cTBS, *v* ~ cTBS, *z* ~ cTBS.

**Model 1 (*a* only):**

*a* ~ cTBS + cTBS:AMB, *v* ~ cTBS, *z* ~ cTBS.

**Model 2 (*v* only):**

*a* ~ cTBS, *v* ~ cTBS + cTBS:AMB, *z* ~ cTBS.

**Model 3 (*z* only):**

*a* ~ cTBS, *v* ~ cTBS, *z* ~ cTBS + cTBS:AMB.

**Model 4 (*a + v*):**

*a* ~ cTBS + cTBS:AMB, *v* ~ cTBS + cTBS:AMB, *z* ~ cTBS.

**Model 5 (*a + z*):**

*a* ~ cTBS + cTBS:AMB, *v* ~ cTBS, *z* ~ cTBS + cTBS:AMB.

**Model 6 (*v + z*):**

*a* ~ cTBS, *v* ~ cTBS + cTBS:AMB, *z* ~ cTBS + cTBS:AMB.

**Model 7 (*a* + *v + z*):**

*a* ~ cTBS + cTBS:AMB, *v* ~ cTBS + cTBS:AMB, *z* ~ cTBS + cTBS:AMB.

We used the deviance information criterion (DIC) to select the winning model. A winning model should have a DIC value that is (1) lower than the baseline model and (2) consistently lower than most of the competing models in all conditions. Supplementary Fig. 7 for comparisons of DIC values across models. Results showed that the model 7 (full model) where cTBS affects all three parameters and all effects interact with stimulus ambiguity had the lowest DIC values in most conditions, and hence it was chosen as the wining model. Moreover, the same as the cTBS effect analyses, we simulated the data based on the estimated parameters, and compared summary statistics between the real and the simulated data. The similarity fell in the 95% credible criteria for full models in all conditions. Therefore, cTBS effects on all three DDM parameters were affected by stimulus ambiguity.

We then did simple main effect analyses to study how cTBS effects on three DDM parameters were modulated by stimulus ambiguity. For each condition, trials were separated into three groups according to the step in the tone/consonant continuum (steps 1 and 5: unambiguous; steps 2 and 4: half ambiguous; step 3: ambiguous). For each group of trials, an independent full model that assumes cTBS affected all three parameters was built and posterior distributions of parameters (compared with sham) were tested. Model and parameter estimations were the same as main effect analyses of TBS effects (see Materials and methods). Results were shown in Supplementary Table 2. Note that for the boundary threshold (*a*) and the drift rate (*v*), significant cTBS effects on parameter distributions were only shown when stimuli were not completely ambiguous (i.e., half or un-ambiguous). Thus, it is possible that when sensory information provides absolutely no valid features for perceptual decision, the dLMC loses its contributions.

### Supplementary figures and tables

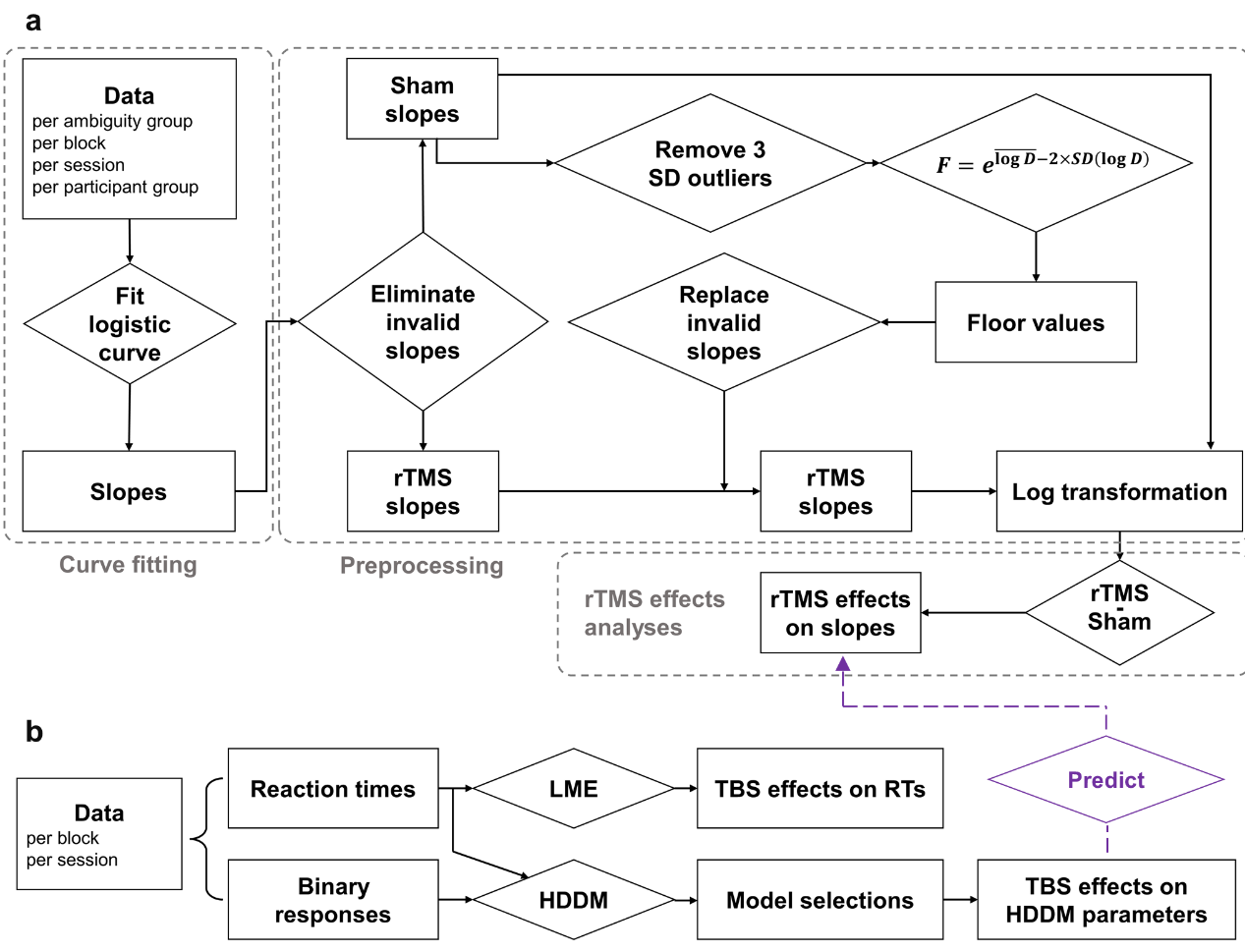

**Supplementary Fig. 1. Data analysis pipelines.**

(A) Perceptual sensitivity analysis includes curve fitting, preprocessing of slopes, and TMS effects analyses (for Experiment 1; those for Experiment 2 were similar).
(B) Latent psychological processes analysis includes modeling, model selections, and post-hoc analyses of TBS effects on model parameters. TBS effects on reaction times were also estimated using linear mixed model estimation.

Besides, we also seek to explain changes in psychometric curve slopes by DDM parameters through comparing results from the two analysis pipelines (purple text).

LME: linear mixed model estimation; HDDM: hierarchical Bayesian estimation of the drift-diffusion model.

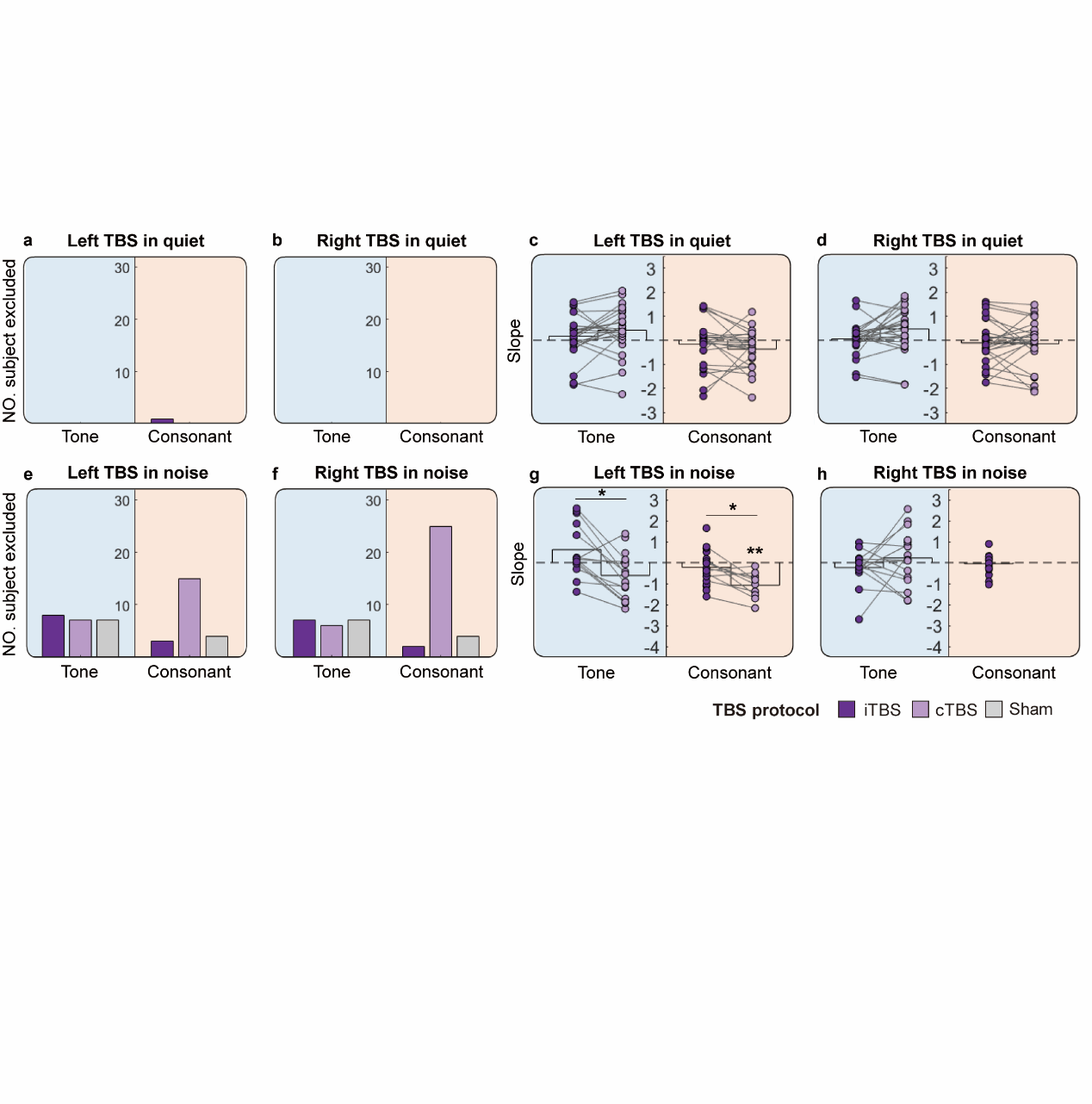

**Supplementary Fig. 2. Elimination of invalid slopes and results without slope replacement in Experiment 2.**

**(a–b) and (e–f)** Number of participants excluded in each condition for whom the psychometric slopes were marked as “invalid” (See Materials and methods, and Supplementary Fig. 1 for details). Note that more invalid results were found in conditions with noise presented, especially for consonant perception in noise (**e–f**).

**(c–d) and (g–h)** Results of psychological curve slopes in Experiment 2 before slope replacement (See Materials and methods, and Supplementary Fig. 1 for details). The same results but with slope replacement are shown in Fig. 3.

See Fig. 2 legend for a detailed description of color patterns.

Statistical tests were performed by non-parametric tests comparing TMS effects with zero, and comparing effects of cTBS with iTBS within the same tasks. P values (single-tailed) were adjusted by false discovery rate (FDR) correction (threshold = 0.05). * *p_fdr_* < 0.05; ** *p_fdr_* < 0.01; *** *p_fdr_* < 0.001.

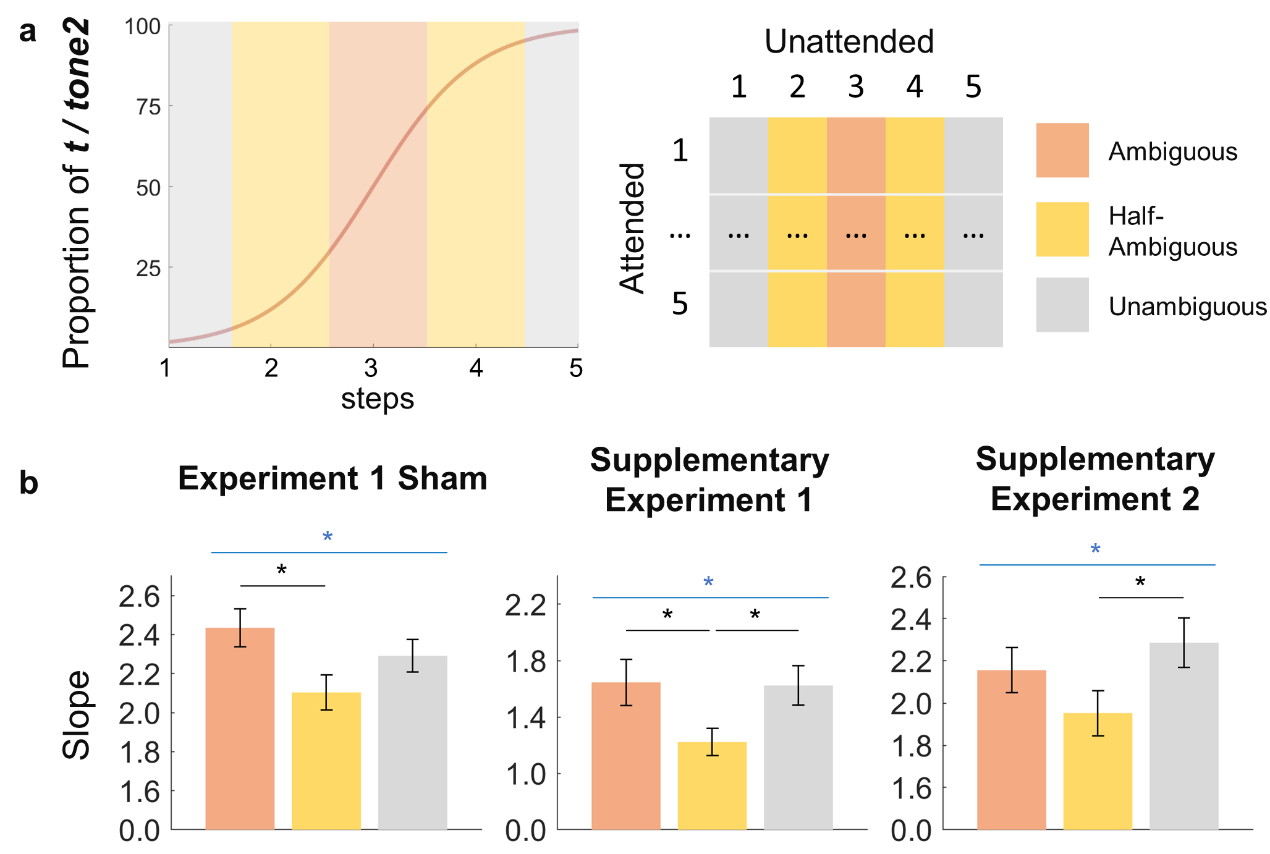

**Supplementary Fig. 3. Competitions between tone and consonant perception.**

(**a**) Take Experiment 1 Sham as an example: trials were grouped according to the ambiguity of the unattended feature. Left: the ambiguity is a function of stimuli continuum (participants made confident judgments at unambiguous steps, biased but uncertain judgements at half-ambiguous steps, and guessing judgments at ambiguous steps). Right: trials were grouped into 3 conditions according to unambiguous, half ambiguous, and ambiguous steps in the unattended dimension. “t” represents [t^h^]; “tone2” represents mid-rising tone. Orange: ambiguous conditions; yellow: half-ambiguous conditions; grey: unambiguous conditions.

(**b**) The ambiguity of the unattended feature affected categorical speech perception. The main effects of ambiguity on slopes of psychometric curves were significant in Experiment 1 Sham (left), Supplementary Experiment 1 (middle), and Supplementary Experiment 2 (right). Post-hoc analyses showed that slopes were significantly lowered in half-ambiguous conditions compared with ambiguous (Experiment 1 Sham and Supplementary Experiment 1) and unambiguous (Supplementary Experiment 1 and Supplementary Experiment 2) conditions. Blue horizontal lines and asterisks represent significant main effects of ambiguity; black horizontal lines and asterisks represent significant differences between ambiguity conditions. Statistical tests were performed by rANOVA for Experiment 1 Sham and Supplementary Experiment 1, and by non-parametric Friedman test and Wilcoxon signed ranks test for Supplementary Experiment 2. P values for post hoc analyses were adjusted by Bonferroni’s multiple comparison corrections. * corrected *p* < 0.05.

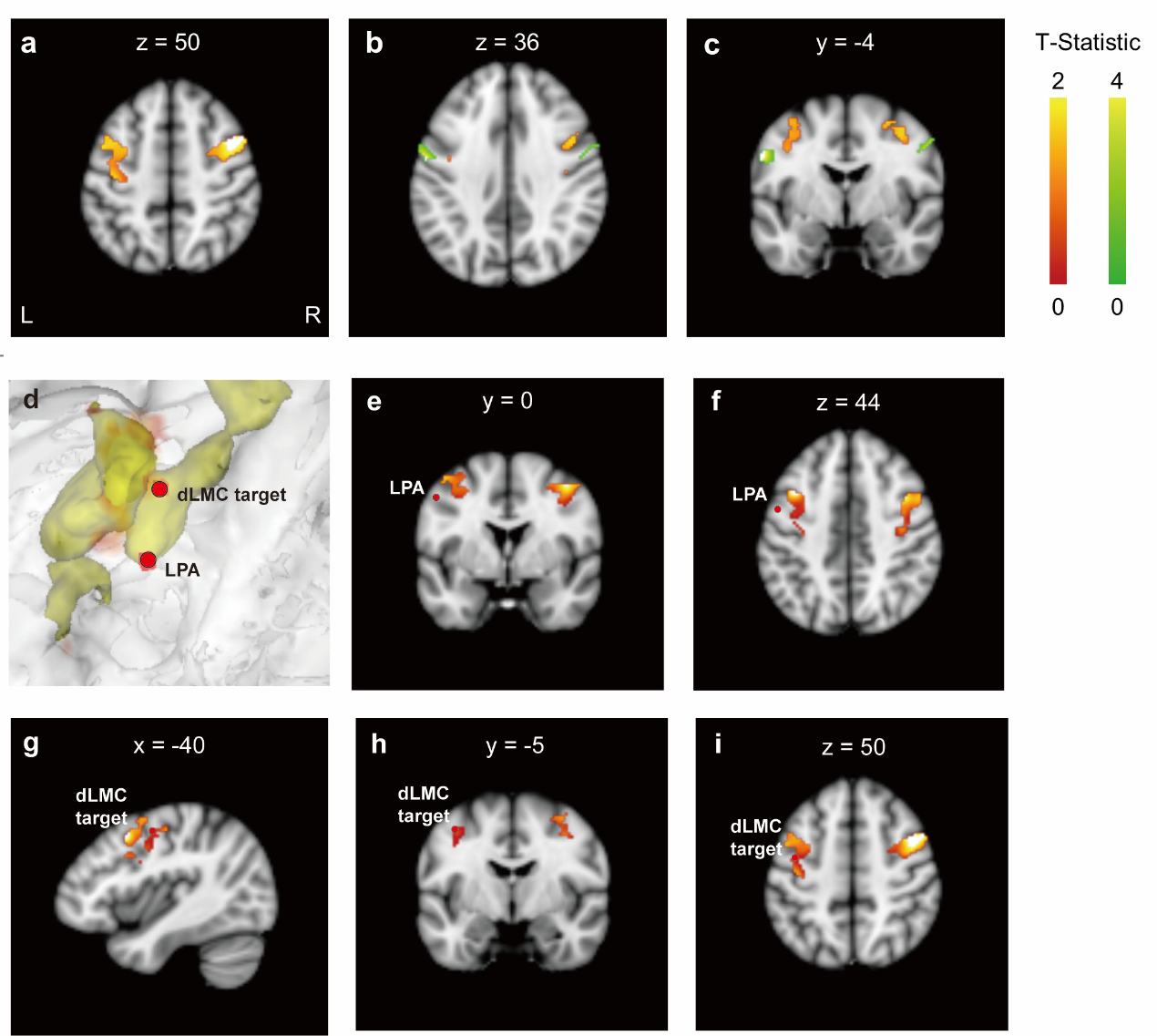

**Supplementary Fig. 4. Group-level activation in the functional localization experiment and the dLMC target location.**

**(a–c)** Group-level activation maps. The axial views show the positions of dLMC (**a and b**, “say AH” – “say D”, orange) and TMC (**b**, “say D” – “say AH”, green) activation areas. **(c)** Dorsal-ventral functional dissociation between larynx (orange) and tongue (green) motor areas is shown from the coronal view of activation.

**(d–i)** Comparison between the location of the used dLMC target in the current study and the activation peak for the dorsal larynx/phonation area (LPA) in Brown et al. (2008)^10^. **(d)** The dLMC target and the dorsal LPA peak on the surface map (red dots) with the activation volumes for dLMC (“say AH” – “say D”, colored areas). Note that, both nodes are within the dorsal-ventral range of the activations. **(e and f)** The spatial relationship between the dorsal LPA (red dot) and the “say AH” – “say D” activation areas (orange areas) from the coronal **(e)** and sagittal **(f)** views. **(g–i)** The spatial relationship between the dLMC target (red dot) and the “say AH” – “say D” activation areas (orange areas) from the axial **(g)**, coronal **(h)**, and sagittal **(i)** views.

Activation areas were masked by AAL ROIs of bilateral precentral gyrus. L, left hemisphere; R, right hemisphere. x, y, and z represent MNI coordinates at the left-right, anterior-posterior, and inferior-superior extents, respectively. Visualization was performed by Mango^13^.

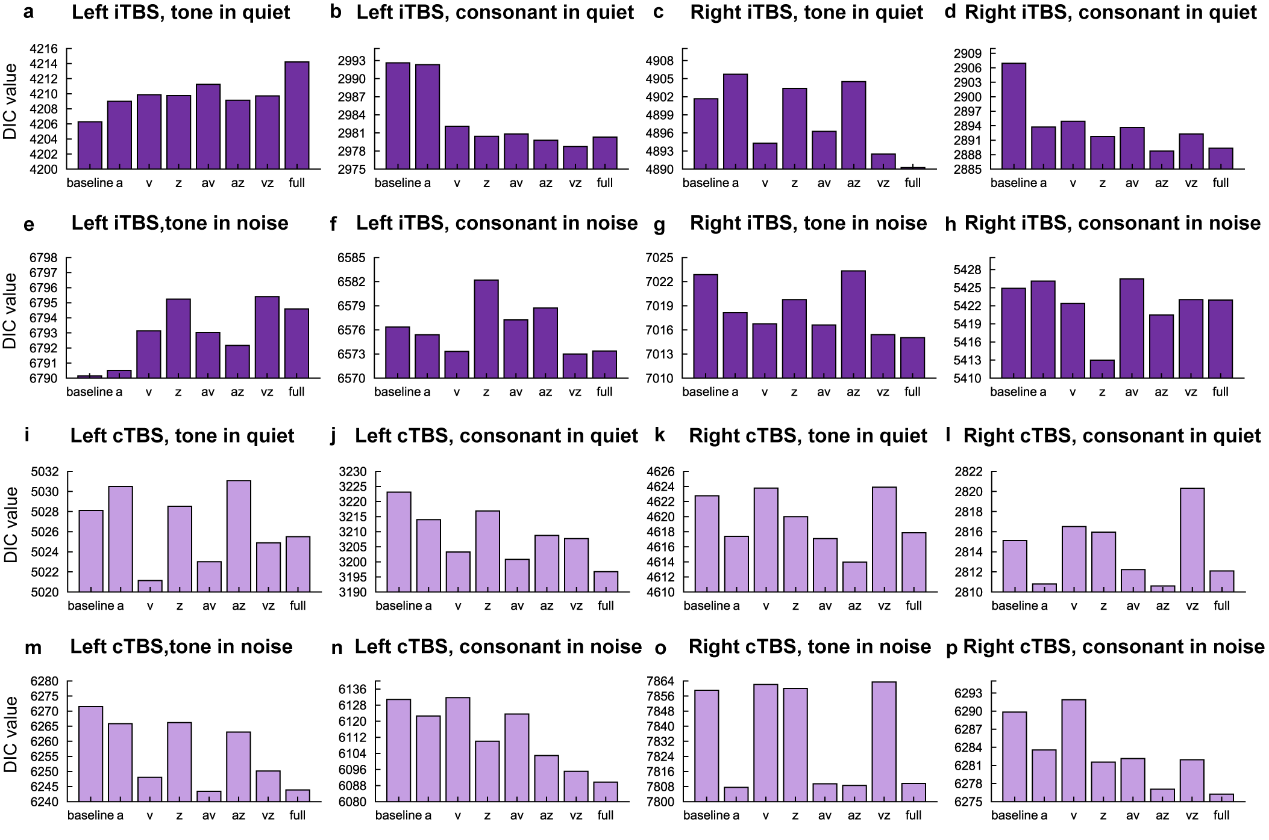

**Supplementary Fig. 5：HDDM deviance information criterion (DIC) values of eight different models in various conditions.**

**(a–h)** Model comparisons in iTBS conditions.

**(i–m)** Model comparisons in cTBS conditions.

DDM parameters: a: threshold for decision boundaries; v: drift rate for evidence accumulation; z: the starting point of evidence accumulation or response bias.

Baseline: baseline regression model (null model) that assumes no TBS effects; a/v/z: models assuming only one of the three DDM parameters is affected by TBS; av/az/vz: models assuming two of the three DDM parameters are affected by TBS; full(avz): the model assuming all three DDM parameters are affected by TBS.

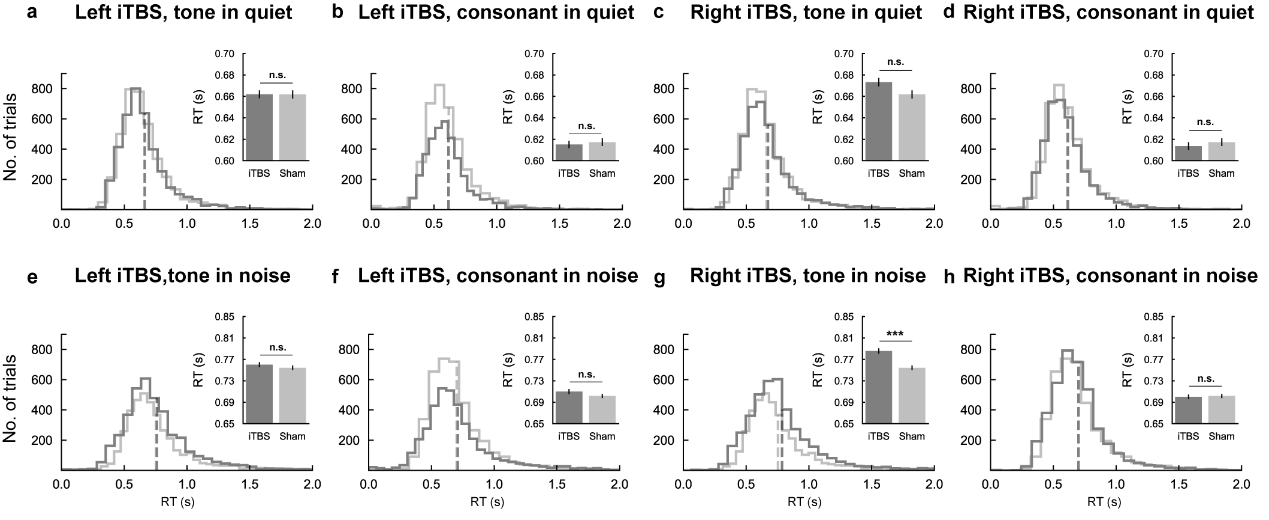

**Supplementary Fig. 6：iTBS effects upon the dLMC on RTs in Experiment 2.**

Compared with sham, iTBS upon the right dLMC increased RTs in the tone perception task in noise (**g**), but did not affect RTs in the other conditions (**a–h**).

For all statistical test, p values (two-tailed) were adjusted by false discovery rate (FDR) correction (threshold = 0.05). * *p_fdr_* < 0.05; ** *p_fdr_* < 0.01; *** *p_fdr_* < 0.001.

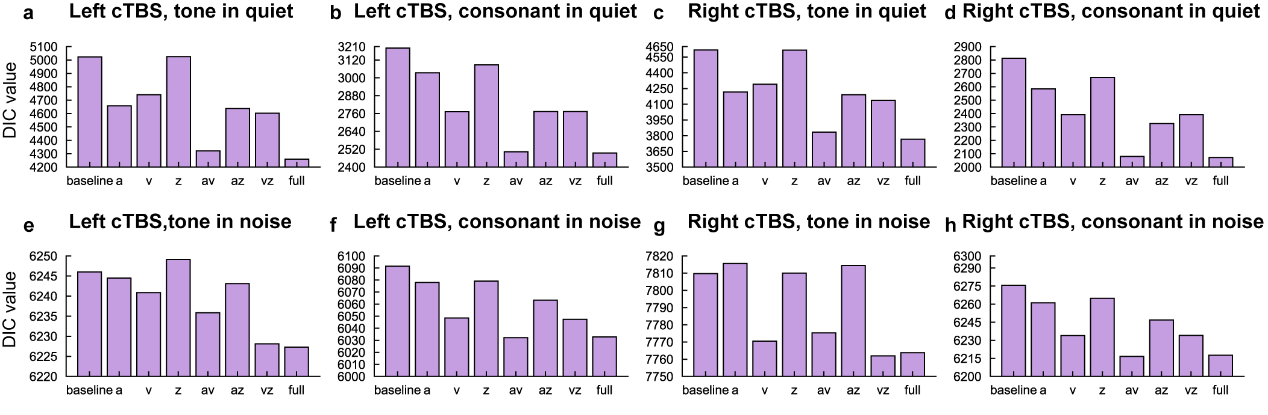

**Supplementary Fig. 7：HDDM deviance information criterion (DIC) values of eight different models with interaction items in various conditions.**

DDM parameters demonstrated are equivalent to Supplementary Fig. 5.

Note that the “baseline” model here is equal to the “full” model that assumes cTBS affects all three parameters but without interactions with stimulus ambiguity; a/v/z: models that assume cTBS affects all three parameters and only one of the effects interacts with stimulus ambiguity; av/az/vz: models that assume cTBS affects all three parameters and two of the effects interact with stimulus ambiguity; full(avz): the model assuming cTBS affects all three parameters and all effects interact with stimulus ambiguity.

**Supplementary Table 1.** **Acoustic properties of stimuli.**

Acoustic properties of clear syllables, and individualized synthesized continua used in Experiment 1 and 2. Note that for Experiment 1, [ti^h^55] – [ti^h^35] and [ti35] – [ti^h^35] were not shown. Values depict mean ± standard deviation. VOT: voice onset time. F0 was extracted by the *pitch* function in MatlabR2021a. F0 glide was calculated by subtracting pitch onsets from pitch offsets.

| **Stimuli** | **VOT (ms)** | **Duration (ms)** | **mean F0 (Hz)** | **F0 glide (Hz)** | **F0 glide (Hz/s)** |
| --- | --- | --- | --- | --- | --- |
| **Clear syllables** | |  |  |  |  |
| **[ti55]** | 0.00 | 355.94 | 334.63 | 7.65 | 27.33 |
| **[ti35]** | 0.00 | 354.06 | 271.39 | 105.75 | 377.68 |
| **[ti^h^55]** | 88.68 | 444.63 | 334.63 | 7.65 | 27.33 |
| **[ti^h^35]** | 88.68 | 442.74 | 271.39 | 105.75 | 377.68 |
| **Experiment 1** | |  |  |  |  |
| **[ti55]**  **\|**  **[ti35]** | 3.41 ± 0.00 | 357.39 ± 0.00 | 333.22 ± 0.06 | 7.62 ± 0.63 | 25.40 ± 2.12 |
|  | 3.41 ± 0.00 | 357.39 ± 0.00 | 327.53 ± 2.74 | 16.01 ± 4.16 | 53.37 ± 13.87 |
|  | 3.41 ± 0.00 | 357.39 ± 0.00 | 321.95 ± 5.46 | 24.81 ± 8.28 | 82.70 ± 27.59 |
|  | 3.41 ± 0.00 | 357.39 ± 0.00 | 316.26 ± 8.08 | 33.10 ± 12.17 | 110.34 ± 40.55 |
|  | 3.41 ± 0.00 | 357.39 ± 0.00 | 310.63 ± 10.81 | 41.43 ± 15.90 | 138.10 ± 53.00 |
| **[ti55]**  **\|**  **[ti^h^55]** | 3.41 ± 0.00 | 357.39 ± 0.00 | 333.22 ± 0.06 | 7.62 ± 0.63 | 25.40 ± 2.12 |
|  | 16.05 ± 2.93 | 370.02 ± 2.93 | 333.22 ± 0.06 | 7.62 ± 0.63 | 25.40 ± 2.12 |
|  | 28.68 ± 5.86 | 382.66 ± 5.86 | 333.22 ± 0.06 | 7.62 ± 0.63 | 25.40 ± 2.12 |
|  | 41.32 ± 8.79 | 395.31 ± 8.79 | 333.22 ± 0.06 | 7.62 ± 0.63 | 25.40 ± 2.12 |
|  | 53.96 ± 11.72 | 407.93 ± 11.72 | 333.22 ± 0.06 | 7.62 ± 0.63 | 25.40 ± 2.12 |
| **Experiment 2** | |  |  |  |  |
| **[ti55]**  **\|**  **[ti35]** | 1.54 ± 1.60 | 355.52 ± 1.60 | 333.58 ± 0.58 | 7.16 ± 0.90 | 23.88 ± 2.99 |
|  | 1.54 ± 1.60 | 355.52 ± 1.60 | 327.63 ± 1.01 | 16.38 ± 1.57 | 54.61 ± 5.23 |
|  | 1.54 ± 1.60 | 355.52 ± 1.60 | 321.72 ± 2.04 | 25.29 ± 3.15 | 84.29 ± 10.48 |
|  | 1.54 ± 1.60 | 355.52 ± 1.60 | 315.82 ± 3.11 | 34.11 ± 4.86 | 113.70 ± 16.19 |
|  | 1.54 ± 1.60 | 355.52 ± 1.60 | 309.93 ± 4.07 | 43.10 ± 6.39 | 143.67 ± 21.31 |
| **[ti55]**  **\|**  **[ti^h^55]** | 1.54 ± 1.60 | 355.52 ± 1.60 | 333.58 ± 0.58 | 7.16 ± 0.90 | 23.88 ± 2.99 |
|  | 11.36 ± 3.10 | 365.24 ± 3.13 | 333.58 ± 0.58 | 7.16 ± 0.90 | 23.88 ± 2.99 |
|  | 21.17 ± 5.49 | 375.13 ± 5.53 | 333.58 ± 0.58 | 7.16 ± 0.90 | 23.88 ± 2.99 |
|  | 30.99 ± 8.00 | 384.95 ± 8.04 | 333.58 ± 0.58 | 7.16 ± 0.90 | 23.88 ± 2.99 |
|  | 40.80 ± 10.56 | 394.78 ± 10.55 | 333.58 ± 0.58 | 7.16 ± 0.90 | 23.88 ± 2.99 |

**Supplementary Table 2. Interactions between effects from cTBS and ambiguity of continua step on DDM parameters.**

Conditions where cTBS exerted significant effects on slopes of psychometric functions are analyzed only. Simple main effect analyses separately in trials with unambiguous, half-ambiguous, and ambiguous tone/VOT are performed if the interaction is significant. +/- signs and numbers represent directions and posterior probabilities of regression coefficients of interactions, or cTBS effects in simple main effect analyses compared with sham (+: higher than sham, -: lower than sham). Probabilities are FDR corrected. Numbers in bold show significant effects (posterior probability p < 0.05, two-tailed). Posterior probability: ’ *p* < 0.1 (marginally significant), * *p* < 0.05, ** *p* < 0.01, *** *p* < 0.001.

|  | **Boundary (a)** | | | | **Drift rate (v)** | | | | **Starting point (z)** | | | |
| --- | --- | --- | --- | --- | --- | --- | --- | --- | --- | --- | --- | --- |
|  | **Inter.** | **UAMB** | **HAMB** | **AMB** | **Inter.** | **UAMB** | **HAMB** | **AMB** | **Inter.** | **UAMB** | **HAMB** | **AMB** |
| **Left cTBS**  **Consonant in quiet** | **+**  **0.000^***^** | +  0.056’ | **+**  **0.048^*^** | 0.576 | **+**  **0.000^***^** | 0.888 | **+**  **0.000^***^** | 0.325 | **+**  **0.016^*^** | 0.244 | 0.444 | 0.6 |
| **Left cTBS**  **Tone in noise** | **+**  **0.006^**^** | 0.243 | 0.308 | 0.576 | **+**  **0.000^***^** | **+**  **0.000^***^** | **+**  **0.000^***^** | 0.860 | **-**  **0.044^*^** | 0.244 | **-**  **0.012^*^** | **+**  **0.036^*^** |
| **Left cTBS**  **Consonant in noise** | **+**  **0.000^***^** | 0.328 | **+**  **0.048^*^** | 0.932 | **+**  **0.000^***^** | **+**  **0.040^*^** | +  0.056’ | 0.325 | -  0.096’ |  |  |  |
| **Right cTBS**  **Consonant in noise** | **+**  **0.000^***^** | +  0.08’ | +  0.056’ | 0.608 | **+**  **0.000^***^** | 0.237 | +  0.056’ | 0.325 | 0.186 |  |  |  |
